## Supplementary figures for "Phylogenomic signatures of repeat-induced point mutations across the fungal kingdom"

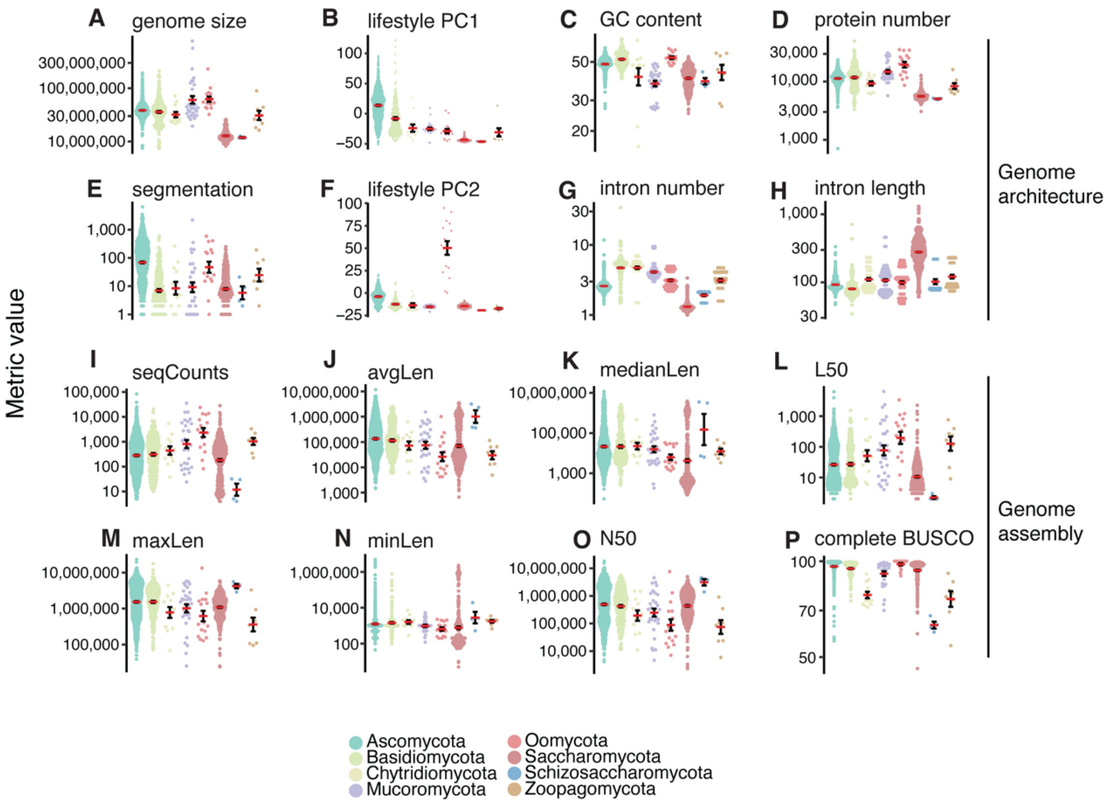


**Fig. S1: Genome assembly metrics across 1,239 fungi. A.** Genome assembly size in total base pairs. **B.** First principal component of the CAZyme repertoire in the genome assembly (lifestyle PC1) suggests that Ascomycetes and Basidiomycetes have an extended repertoire compared to other phyla **C.** GC content of the genome assembly in percentage **D.** Number of annotated protein-coding genes in the assembly **E.** Number of segments with contrasted GC dinucleotide content across the assembly (segmentation) **F.** Second principal component of the CAZyme repertoire in the genome assembly (lifestyle PC2) **G.** Average number of introns per gene in each assembly **H.** Average intron length (bp) in each assembly **I.** Total sequence counts in the assembly (seqCounts) **J.** Average sequence length in the assembly (avgLen) **K.** Median sequence length in the assembly (medianLen) **L.** Count of smallest number of contigs whose length sum makes up half of assembly size (L50) **M.** Maximum sequence length in the assembly (maxLen) **N.** Minimum sequence length in the assembly (minLen) **O.** Sequence length of the shortest contig at 50% of the total assembly length (N50) **P.** Percentage of BUSCO genes found to be complete in each assembly. Note that Oomycota and Mucoromycota have on average larger genomes and more genes while yeasts in the Saccharomycotina subphylum have smaller genomes and less genes. Intron numbers per gene and GC content greatly varies across the dataset, with yeast species having fewer but longer introns and low GC content compared to other Ascomycetes.

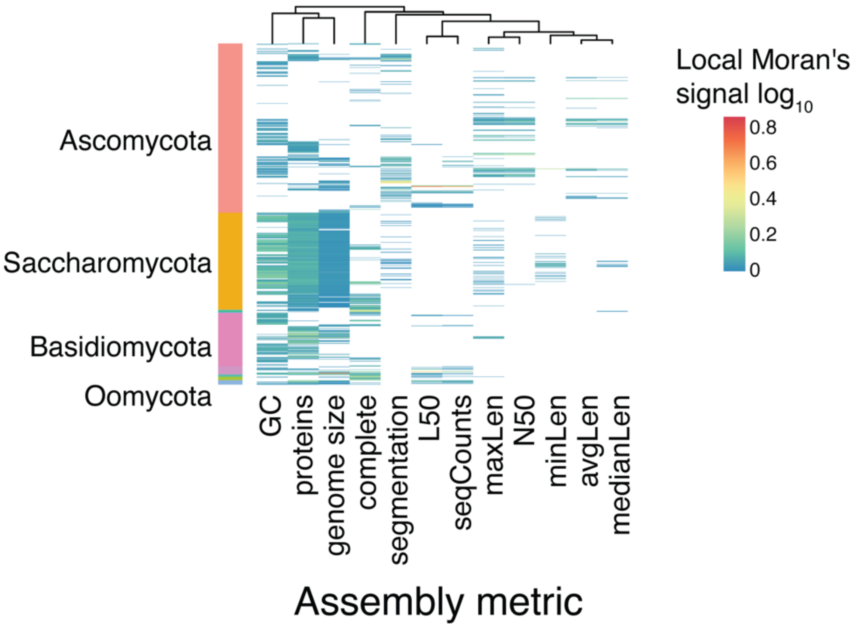


**Fig. S2: Widespread signatures of phylogenetic signal of genome assembly metrics.** Local Indicator of Phylogenetic Association (local Moran's I, log10 transformed for display) for each genome assembly metric as calculated by the *lipaMoran* function from the *phylosignal* R package. Only significant associations are displayed (p-value < 0.05 based on 1,000 permutations). Phylogenetic signal was inferred using five different statistics, namely, Blomberg's K and K*, Abouheif's Cmean, Moran's I, and Pagel's Lambda). Note that two species in the Mucoromycota with some of the largest genomes in the dataset show strong phylogenetic signal for genome size, L50 and the number of scaffolds (Supplementary Table S4, seqCounts).

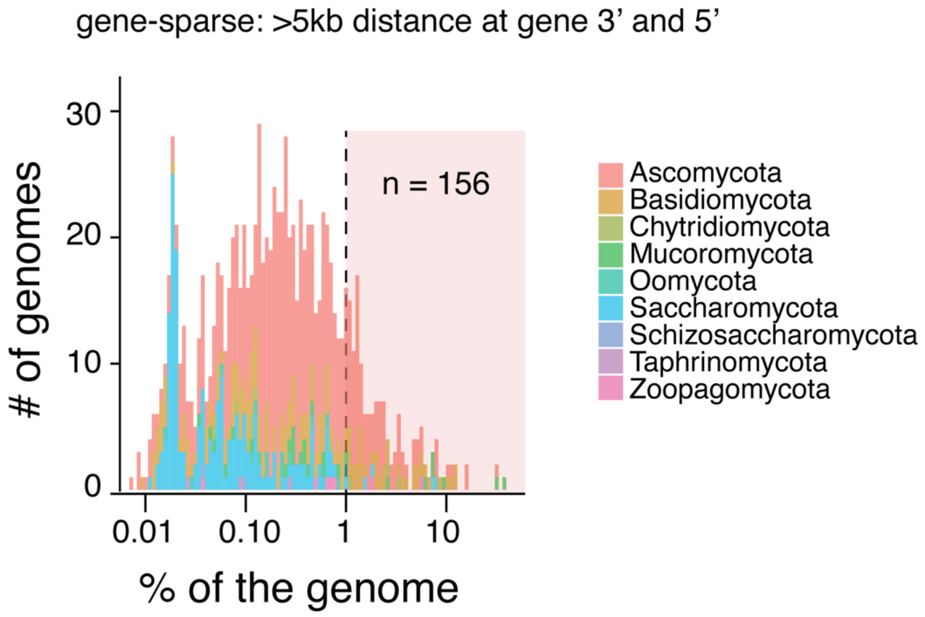


**Fig. S3: Estimation of the number of assemblies showing a "two-speed"-like genome architecture.** Percentage of the gene pool per genome that is found in gene-sparse regions as estimated by >5 kb intergenic distances up and downstream of the gene. Genome assemblies with more than 1% of their gene pool (*n*=156) in such gene-sparse regions are matching a "two-speed" genome architecture.

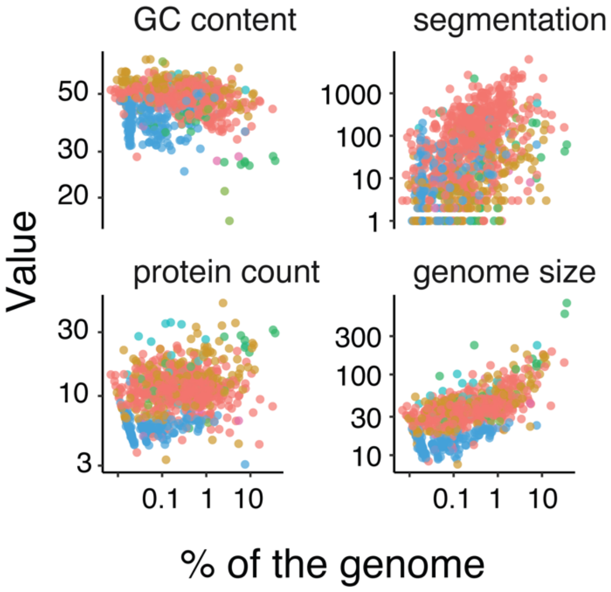


**Fig. S3: Genome compartmentalization strongly correlates with genome size.** Correlation of the percentage of the gene-pool (genome) that is found in gene-sparse regions as estimated by more than 5 kb intergenic distances up and downstream of the gene with genome assembly GC content (%), the number of segments with contrasted GC dinucleotide content across the assembly (segmentation), total protein number and genome assembly size (Mb).

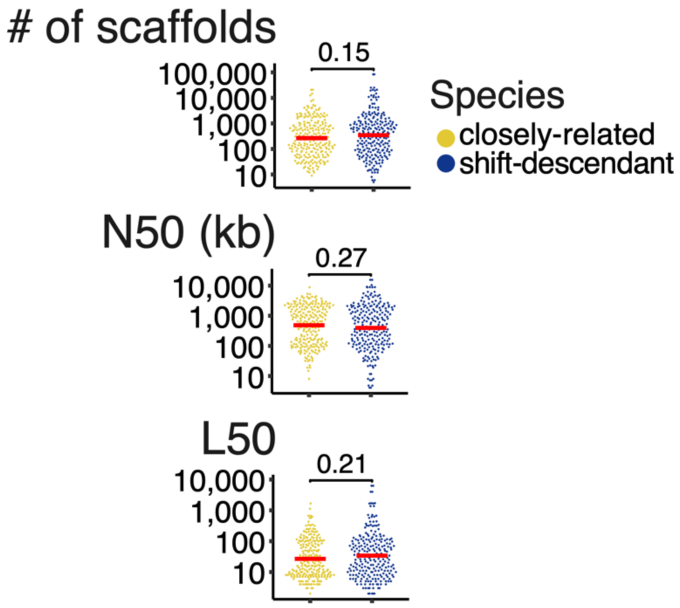


**S5 Fig. No major difference in genome assembly quality between species with an associated shift in genome architecture and their close-relative.** Genome assembly quality given the number of scaffolds, N50 and L50 values for species associated with at a shift in one metric of genome architecture (shift-descendent) compared to close relatives with no shift (closely-related).

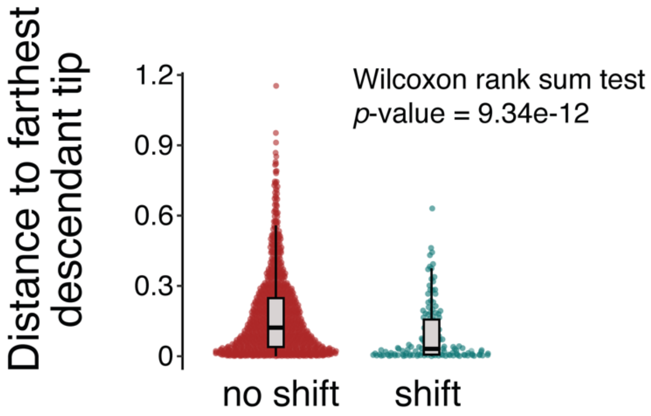


**Fig. S6: Detection of shifts and phylogenetic distances to farthest descendant tip.**

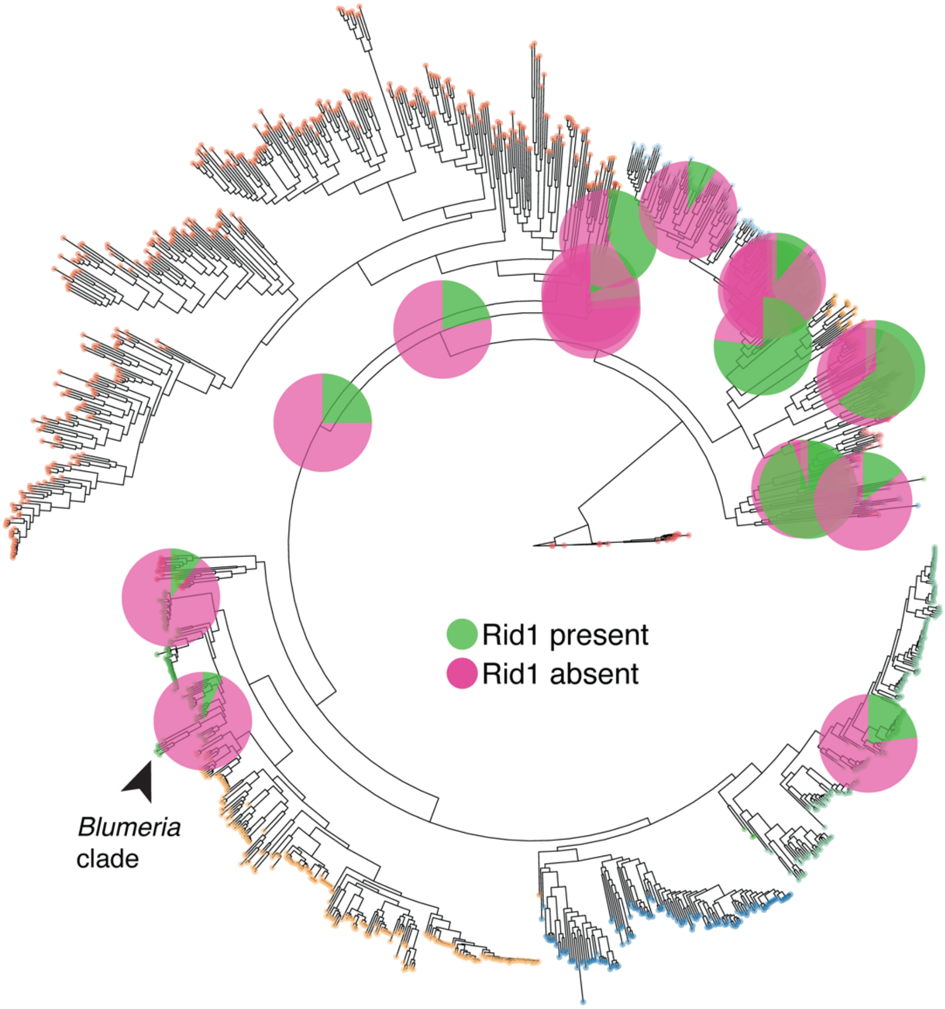


**Fig. S7: Ancestral state reconstruction of presence / absence of the DNA methylase Rid1 across nodes of the phylogeny.** Ancestral state reconstruction was performed using the *ace* function (*ape* package version 5.8 in R), upon the all-rates-different maximum likelihood model of discrete trait. Only nodes with a minimal estimated status probability > 5% are shown.

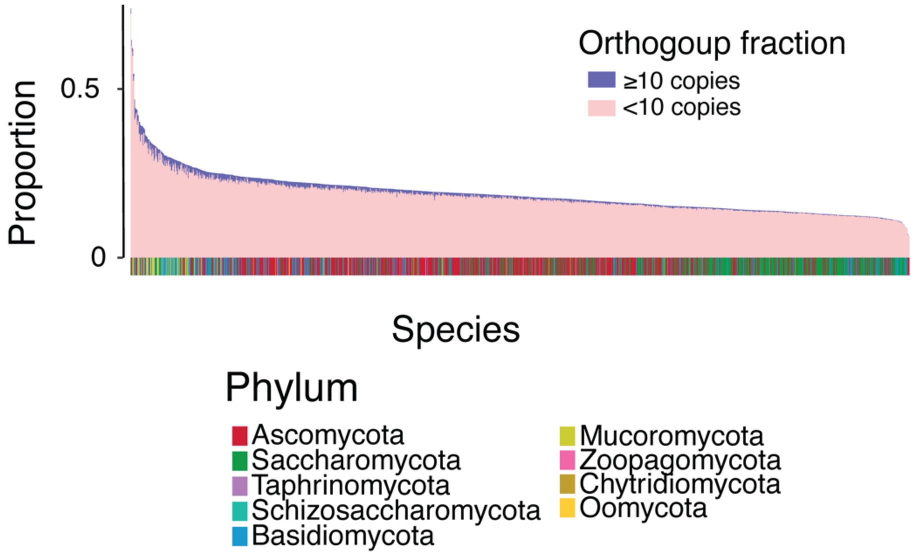


**Fig. S8: Proportion of multi-copy orthogroups identified in each genome.** For each genome, the proportion of orthogroups with counts <10 and >10 protein-coding genes are shown separately.

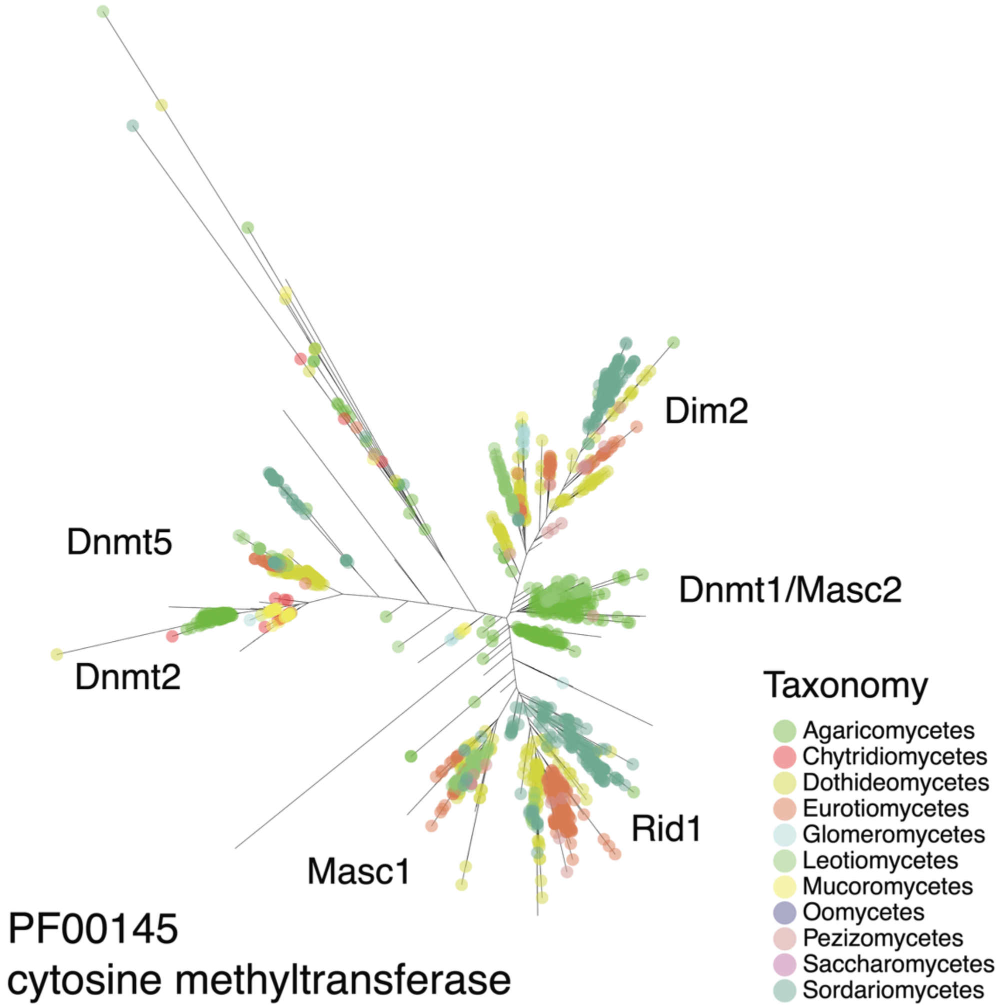


**Fig. S9: Phylogenetic relationship of proteins with a DNA methyltransferase domain (PF00145).** Proteins with an HMM hit to the PF00145 domain were aligned with Clustal Omega and gaps removed using trimAl for phylogenetic reconstruction under the Whelan and Goldman model of protein evolution with fastTree.

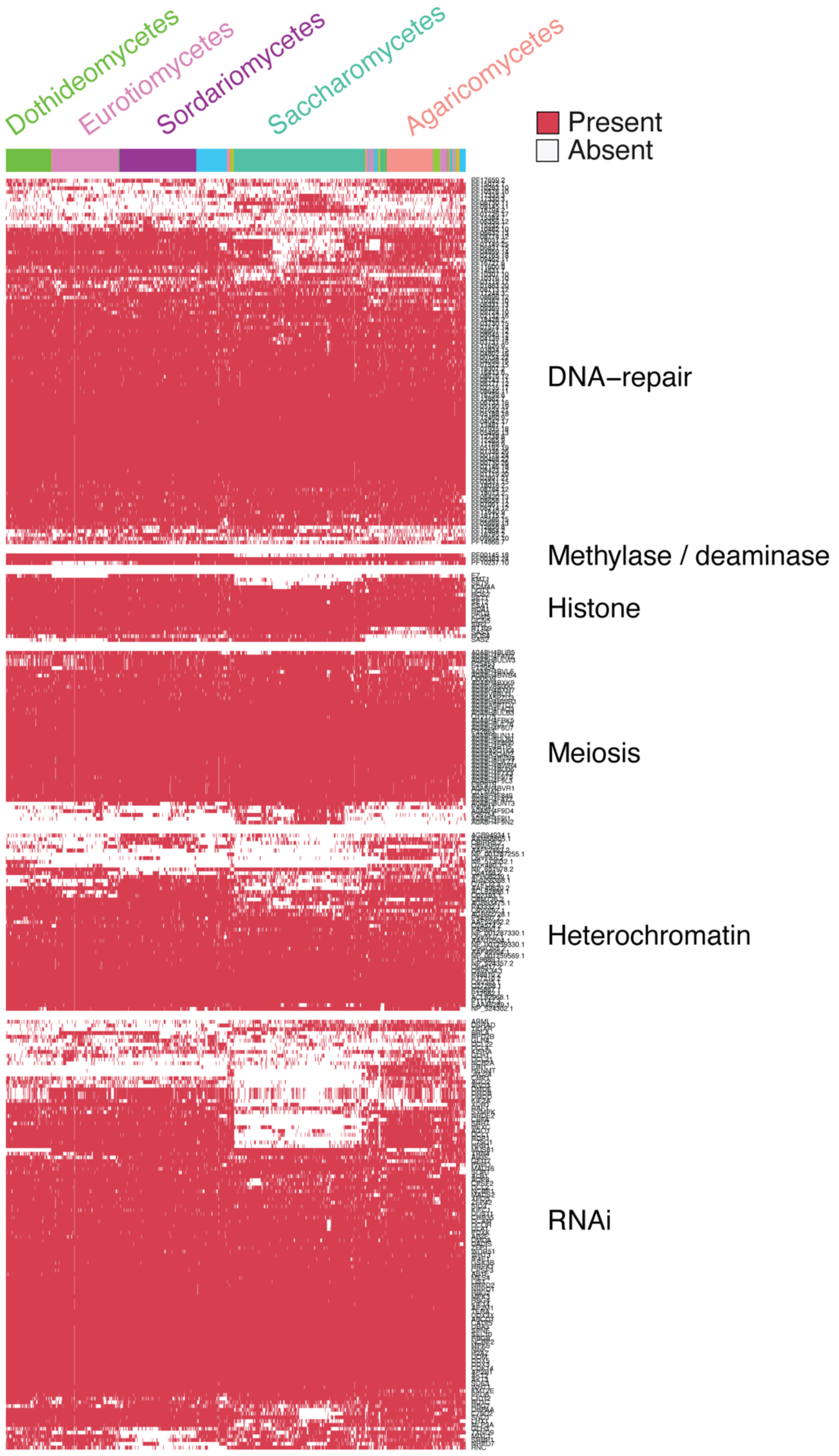


**Fig. S10: Presence / absence heatmap of DNA biology related genes.** For each genome, the presence of functional domains (Pfam) or protein-coding genes known to be involved in DNA repair, DNA methylation, histone biology, meiotic recombination, heterochromatin formation, and RNA interference was assessed by HMM domain or blast search (Supplementary Table S10). Most of the species from the Eurotiomycetes and Dothideomycetes lack the SAS2 Histone acetyltransferase, while species from the Sordariomycetes mostly lack the histone deacetylase 3 (Fig. 4D, SAS2 and NP_651978.2). Most recombination-related genes are conserved in fungi, with the known exceptions of the Mus81 endonuclease absent from Dothideomycetes and the Hop2-Mnd1 recombination complex that is nearly absent from Sordariomycetes. The replication checkpoint Rad17 and the sporulation protein Spo22 are nearly exclusive to the Saccharomycota clade. We also find that homologs of the meiotic sister chromatid recombination protein 1 (Msc1) are absent from the Oomycota, Chytridiomycota, Mucoromycota and Zoopagomycota. In addition to Msc1, Oomycota also lack Msc7 homologs. Similarly, the DNA helicase Srs-2 has been lost multiple times in fungi, including in the Eurotiomycetes. As opposed to the cytosine methyltransferase protein family, which is mostly conserved in Ascomycetes (excluding Saccharomycetes), the adenine methylation domain is conserved in the Saccharomycota but nearly absent in species from the Eurotiomycetes (Fig. 4D, PF10237.10). Protein families related to the double-strand recombination repair are mostly missing in species from the Sordariomycetes (Fig. 4D, PF10376.10). Similarly, proteins from the DNA-mismatch repair family are mostly conserved in Ascomycota but nearly absent from the Eurotiomycetes (Fig. 4D, PF18795.2). Proteins with a PBZ domain, often associated with DNA strand-break repair, are nearly absent from the Ascomycota except for species from the Eurotiomycetes (Fig. 4D, PF10283.10).

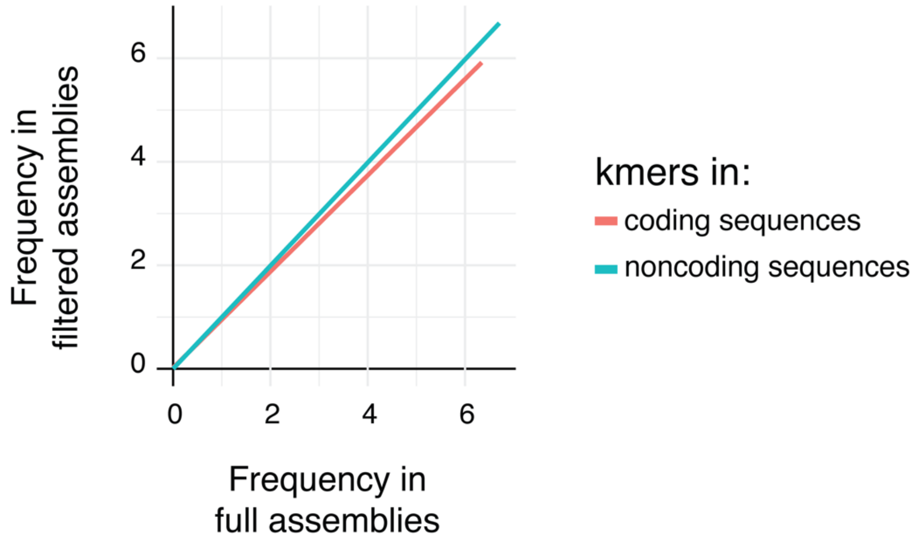


**Fig. S11: Impact of assembly scaffold filtering on *k-mer* frequency at coding and non-coding sequences.** Assemblies were filtered for scaffolds larger than 50 kb (*i.e.* filtered assemblies) and *k-mer* frequency calculated at coding and non-coding sequences (2-, 3- and 4-mers). Although the overall distribution of the frequency values in non-coding sequences is constant across the full and the 50-kb filtered datasets, we find that high frequency *k-mers* tend to be over-represented in coding sequences (1,235 genomes left after filtering due to low assembly contiguity).

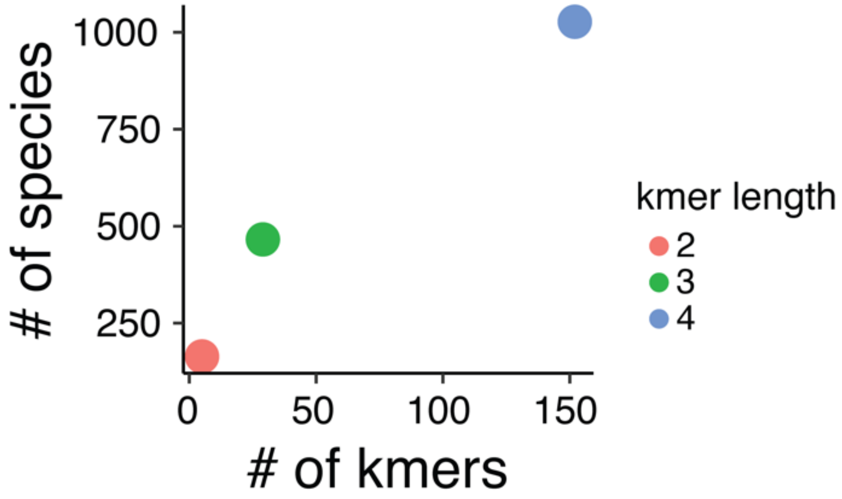


**Fig. S12: Number of >2-fold overrepresented *k-mers* in non-coding sequences compared to coding sequences.** The y-axis shows the number of species in which a 2-, 3-, or 4-mers is found >2-fold overrepresented in non-coding sequences. For the 212 species without a single *k-mer* overrepresented in the non-coding compartment, we find at least one representative of all nine phyla or sub-phyla in the dataset.

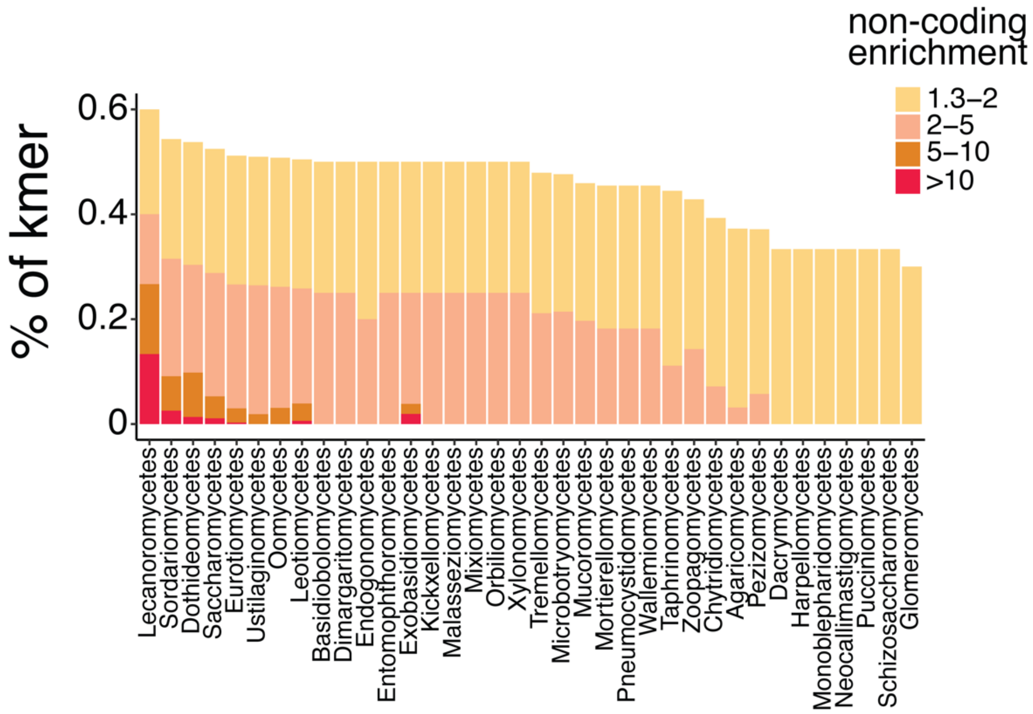


**Fig. S13: Intensity of non-coding *k-mer* enrichment across different taxonomic classes.** Proportion of *k-mer* in each taxonomic class that are enriched at non-coding sequences in the range of 1.3 to 2-fold, 2 to 5-fold, 5 to 10-fold or more than 10-fold. Most taxonomic classes have at least one *k-mer* >2-fold enriched in non-coding sequences (80% or 29 / 36), with Sordariomycetes, Saccharomycetes and Dothideomycetes being the most represented classes, in addition to Eurotiomycetes, Exobasidiomycetes, Lecanoromycetes and Leotiomycetes.

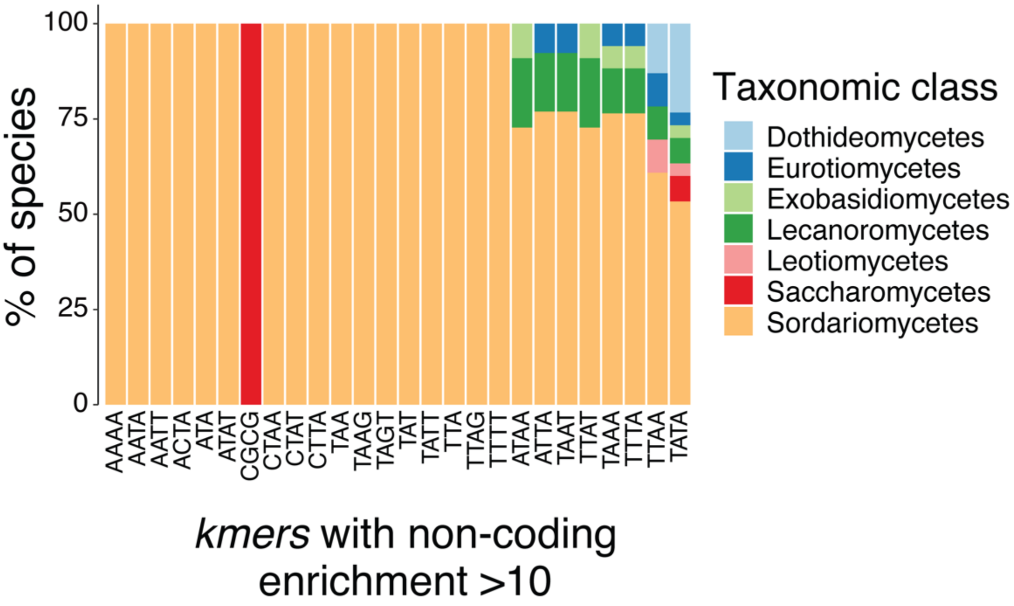


**Fig. S14: Taxonomic distribution of highly enriched *k-mers* at non-coding sequences.** Distribution of the species taxonomic classes carrying the respective *k-mer* more than 10-fold enriched in non-coding sequences.

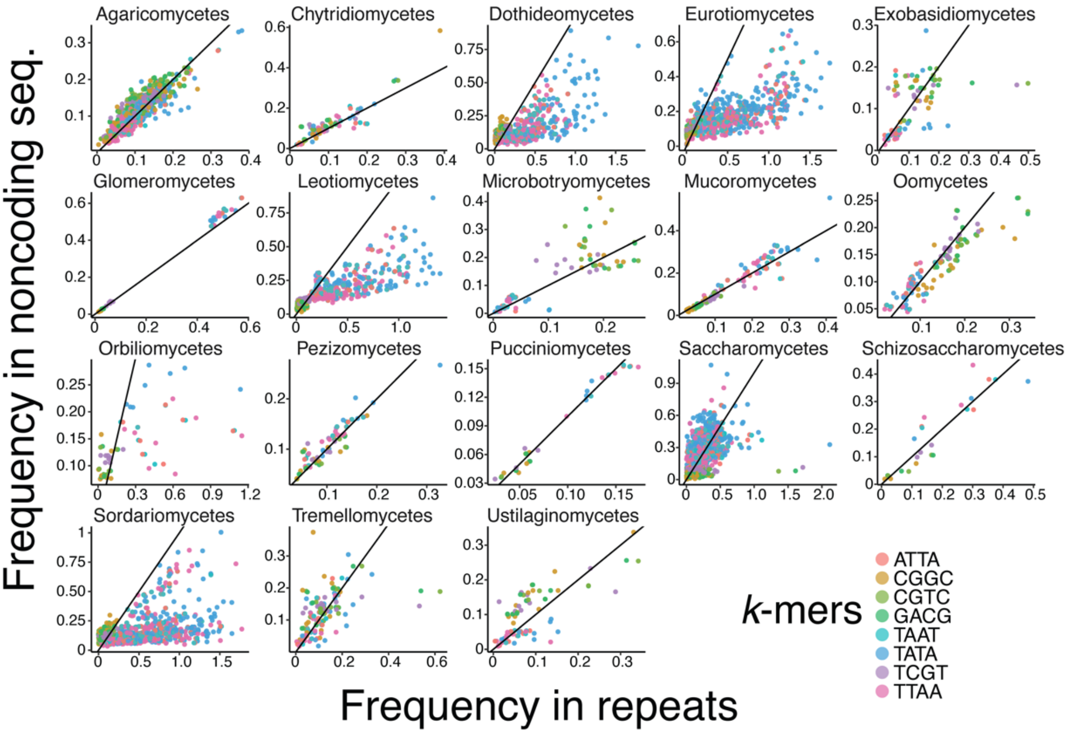


**Fig. S15: *K-mer* frequency at repeats and non-coding sequences across taxonomic classes.** Frequency of eight highly enriched *k-mer* at repeats versus non-coding sequences. Lines represent the y = x diagonal.

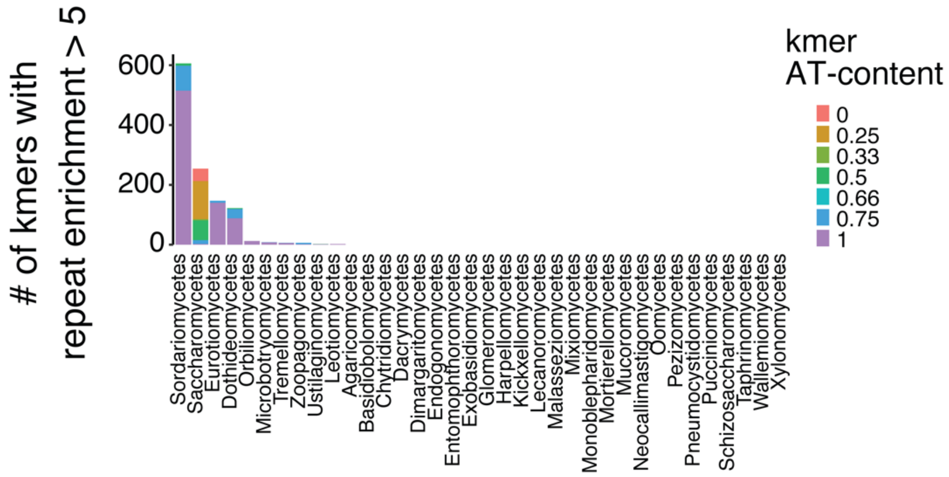


**Fig. S16: Number of *k-mers* with repeat enrichment >5-fold across taxonomic classes.** Total number of *k-mers* with repeat frequency >5-fold compared to non-coding sequences for all 36 taxonomic classes. *K-mers* were split according to their AT-content (0, 0.25, 0.33, 0.50, 0.66, 0.75 or 1).

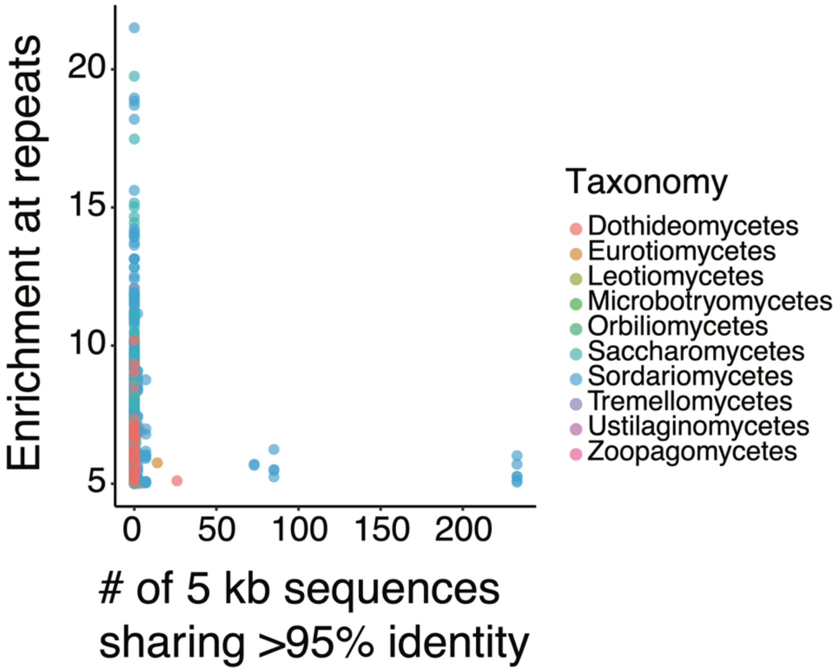


**Fig. S17: Strong enrichment of *k-mers* at repeats associate with few highly similar repetitive sequences.** Genomes with a value of *k-mer* enrichment at repeats >5-fold as a function of the total number of sequences larger than 5 kb sharing between 95-100% sequence identity. Dot colors indicate different taxonomic classes.

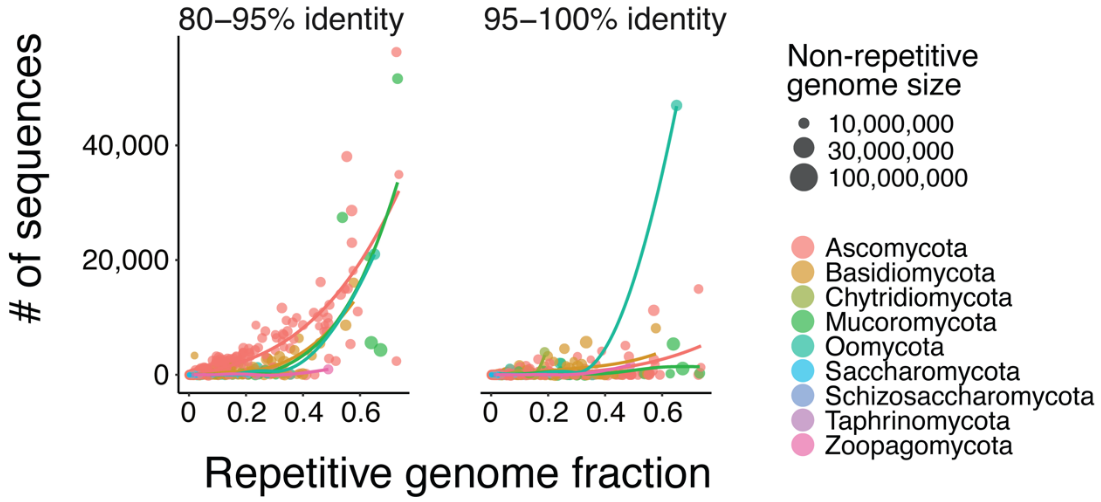


**Fig. S18: High number of highly similar repeats correlates with genome repetitive fraction.** Total number of sequences larger than 5 kb sharing between 80-95% or 95-100% sequence identity per genome assembly as a function of the fraction of the genome assembly identified as repeats. Dot size is scaled to the size of the non-repetitive genome assembly (in base pairs).

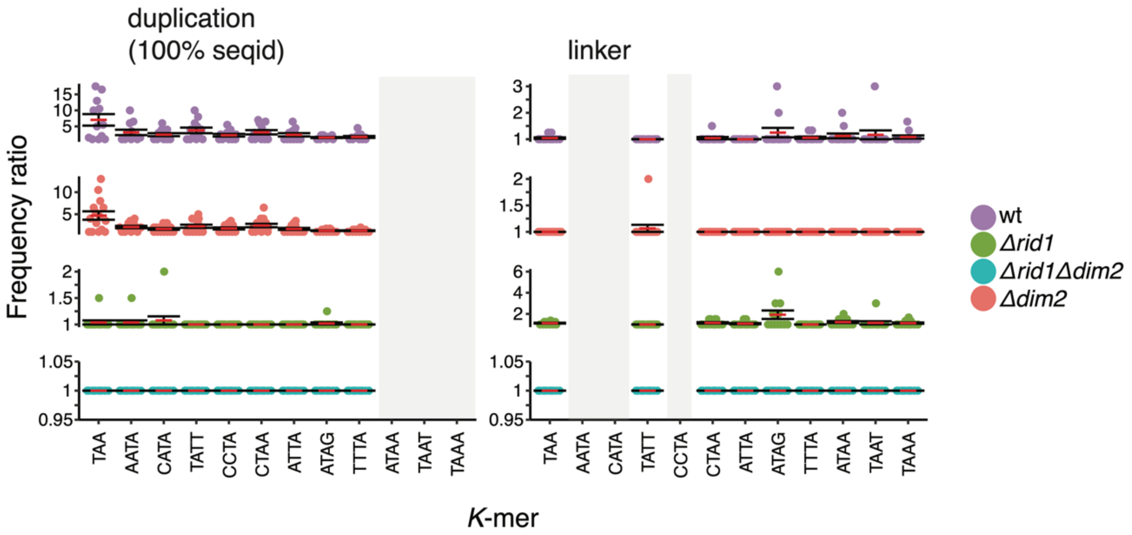


**Fig. S19: Changes in *k-mer* frequency induced by RIP upon a single cross in *Neurospora crassa*.** Frequency ratios of 12 *k-mers* showing >5-fold enrichment at the RLR sequence in the progeny after crosses with the wild-type parents or the deletion mutants *Δdim2*, *Δrid1* and *Δrid1Δdim2*. *K-mer* frequencies were calculated at the duplicated (100% sequence identity) and the unique linker region separately.

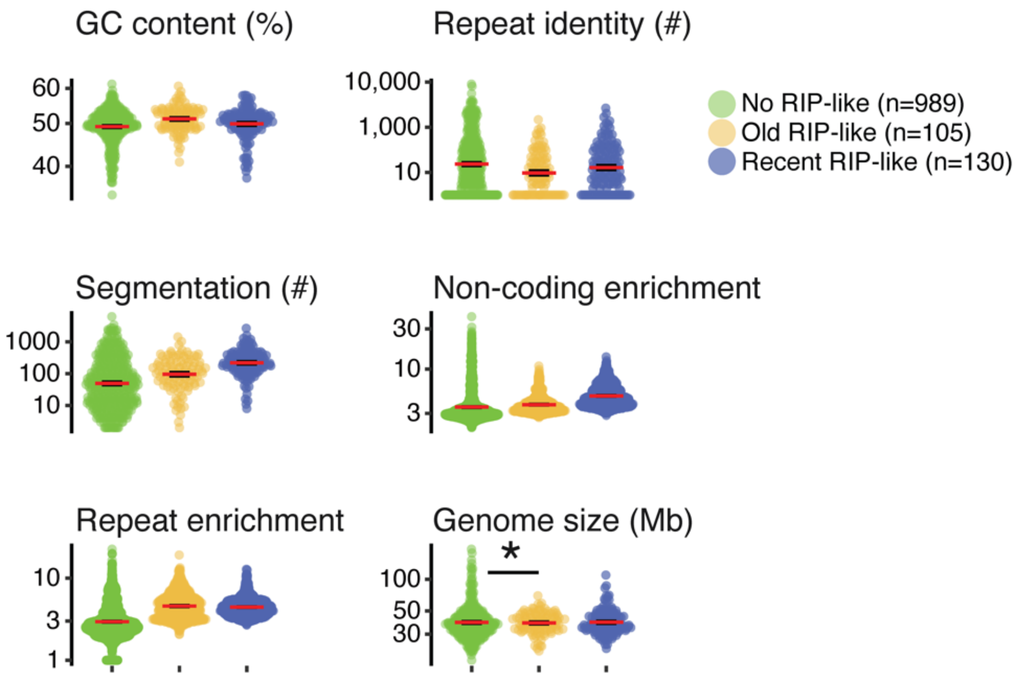


**Fig. S20: Genome architecture in species with evidence for recent or old RIP mutational signatures.** The asterisk denotes TukeyHSD post-hoc test significant differences.

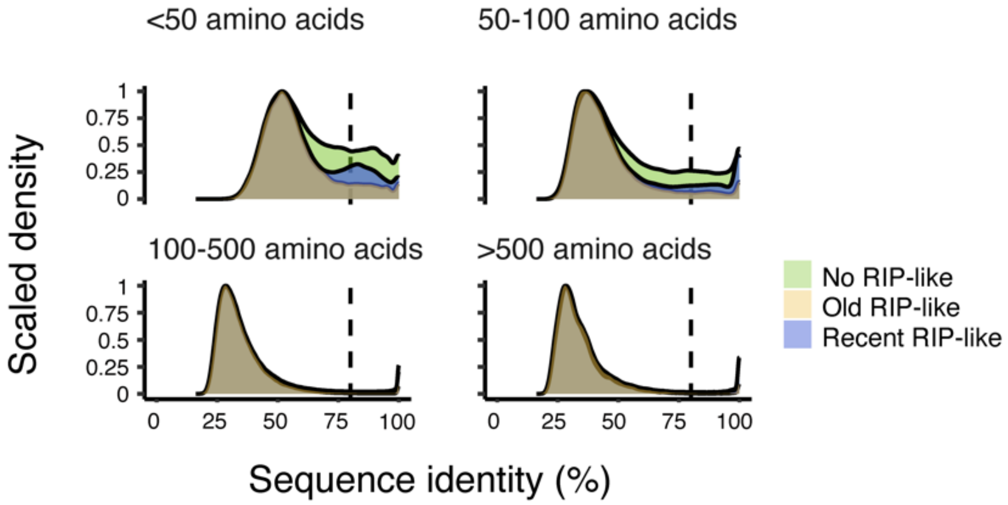


**Fig. S21: Species with no evidence for RIP mutation signatures carry more proteins of high sequence identity.** The x-axis shows protein percentage sequence identity calculated from reciprocal blasts. The y-axis is the scaled density given the number of blast hits. Blast hits with sequence length < 50, between 50 and 100, between 100 and 500 or larger than 500 amino acids are represented in the four facets.

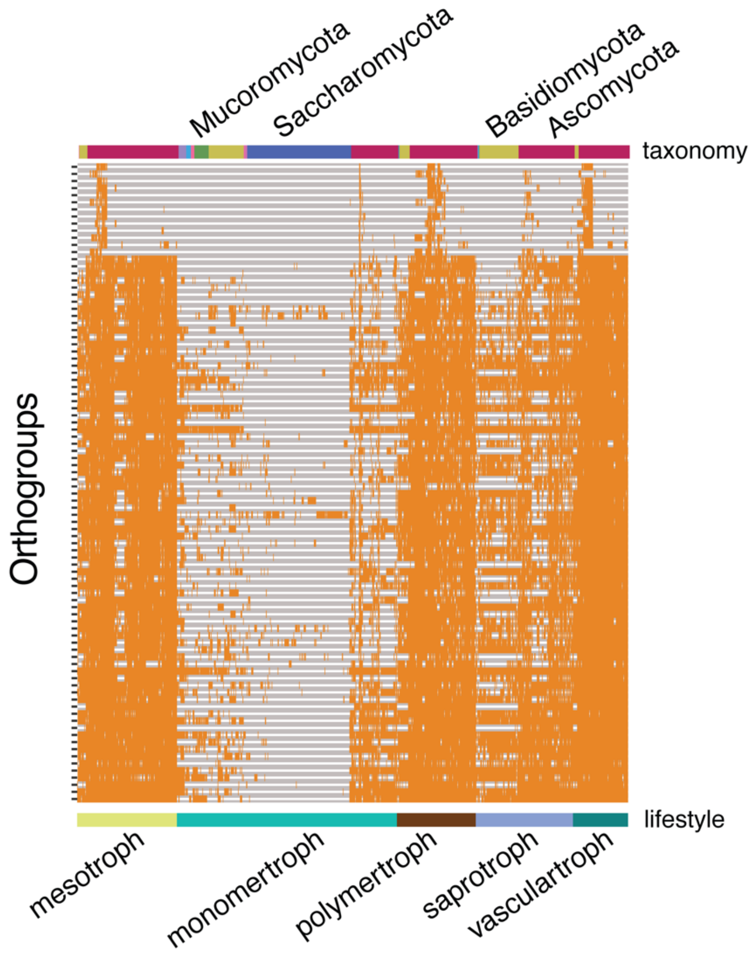


**Fig. S22: Presence / absence heatmap of orthogroups associated with genome size, *k-mer* repeat enrichment and segmentation.** For each associated orthogroup, presence in the focal genome assembly is represented by filled boxes, with the species corresponding taxonomy and lifestyle annotated as colored tiles (top and bottom tiles, respectively).

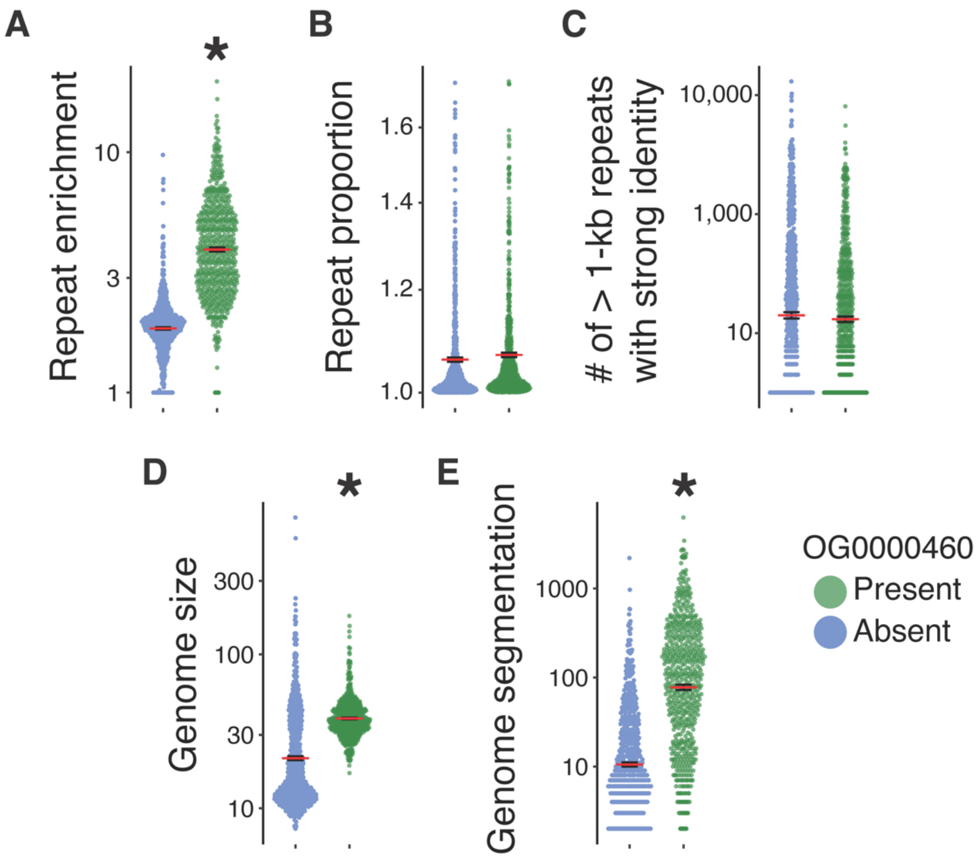


**Fig. S23: Genome architecture in species with and without the repeat enrichment-associated orthogroup OG0000460. A.** TATA *k-mer* enrichment at repeats compared to non-coding sequences in species with or without the OG0000460. **B.** Proportion of the genome annotated as repeats in species with or without the OG0000460 **C.** Number of highly similar (>95% sequence identity) repeated sequences larger than 1 kb in species with or without the OG0000460 **D.** Genome size in species with or without the OG0000460 **E.** Number of isochore genome segments in species with or without the OG0000460. Asterisks denote significant difference as per a TukeyHSD post-hoc test.
